# LiCl-induced sickness modulates spontaneous activity and response dynamics in rat gustatory cortex

**DOI:** 10.1101/2022.01.13.476147

**Authors:** Bradly T. Stone, Jian-You Lin, Abuzar Mahmood, Alden J. Sanford, Donald B. Katz

**Author notes:** <u>Corresponding Author</u> Don Katz, Department of Psychology/Neuroscience Program. <u>Affiliation Postal Code</u> Brandeis University, 415 South Street, Waltham, MA 02453.

## Abstract

Gustatory Cortex (GC), a structure deeply involved in the making of consumption decisions, presumably performs this function by integrating information about taste, experiences, and internal states related to the animal’s health, such as illness. Here, we investigated this assertion, examining whether illness is represented in GC activity, and how this representation impacts taste responses and behavior. We recorded GC single-neuron activity and local field potentials (LFP) from healthy rats and (the same) rats made ill (*via* LiCl injection). We show (consistent with the extant literature) that the onset of illness-related behaviors arises contemporaneously with alterations in spontaneous 7-12Hz LFP power at ∼11 min following injection. This process was accompanied by reductions in single-neuron taste response magnitudes and discriminability, and with enhancements in palatability-relatedness – a result reflecting the collapse of responses toward a simple “good-bad” code arising in a specific subset of GC neurons. Overall, our data show that a state (illness) that profoundly reduces consumption changes basic properties of the sensory cortical response to tastes, in a manner that can easily explain illness’ impact on consumption.

**Significance Statement:** Neural responses in primary sensory cortex are often thought to faithfully represent physical stimuli, and while recent studies (including ours) have challenged this view by documenting enhancements and decrements in stimulus-induced firing related to animals’ internal states, there has been little work setting these changes in any sort of functional, mechanistic context. Here we show that a state (illness) that profoundly reduces consumption changes basic properties of the sensory cortical response to tastes, and then go beyond this to precisely characterize this response plasticity, connecting it to the specific perceptual changes that drive illness’ impact on consumption.

## Introduction

A host of external and internal contextual variables work with experience to shape an animal’s behavior in response to sensory stimulation. Prominent among these variables are the animal’s own internal states, which can be extremely positive (e.g., “euphoria”) or negative (e.g., “depression”), and which profoundly influence the animal’s interactions with its environment [1]; systemic illness, for instance, such as that induced by the intake of toxins, drastically alters an animal’s behavior in relation to food stimuli [2-4]. Such states are likely instantiated in broadly distributed neural networks, and as such, they can be indexed using spectral properties of the electroencephalogram [5, 6] or local field potentials (LFP; [7, 8])

One mechanism whereby illness might influence feeding behavior is *via* modifications of taste perception. Sickness changes taste palatability [9, 10], making even normally preferred substances aversive (potentially prolonging infirmity by demotivating the ingestion of nutrients and/or a possible cure for said illness [11]). This impact of illness on taste palatability can be long-lasting, and even permanent [12], an intimacy of interaction that makes it reasonable to propose that illness may manifest, at least in part, as changes in the function of brain regions within which one can record palatability-related taste responses.

One such region is gustatory cortex (GC), which is involved both in processing of palatability and illness-related learning [13, 14]. As of now, however, little is known about the cortical processing of malaise. Much is known about the basic mechanisms of function for malaise-causing agents such as lithium chloride (LiCl) at the non-neural and peripheral levels [2-4, 15-19], but the impact of these basic mechanisms on activity in regions critical for chemosensory learning remains uninvestigated.

The work presented here begins to fill that knowledge gap, taking a cue from studies showing that basic information about brain states can be assessed in analysis of LFPs, and more specifically from studies showing: 1) that spectral properties of LFPs change with changes in an animal’s internal state [20, 21]; and 2) that these changes are coupled with changes in single-neuron firing dynamics [22-24]. These effects apply to GC taste coding [25], which change with changes in attentional state in a manner that is specifically linked to palatability coding [26]. Here, using extracellular single-neuron recordings, we test the degree to which the onset of an illness state (indexed in terms of changes in mobility) is related to changes in GC LFPs, and go on to test how these behavioral and LFP phenomena correlate with changes in identity-related and hedonic information in GC single-neuron ensemble responses.

Our results demonstrate that the illness-induced changes in GC *μ*(7-12Hz) power reflect the onset (but not the maintenance) of sickness, emerging around the first appearance of sickness-related behaviors [2-4, 27, 28] but subsiding after the transition to the illness state. This change is accompanied by reductions in the magnitude of taste-driven responses, which convey less information about taste identity in sick rats, and simultaneously by enhancement in the palatability-relatedness of the same responses, which appear to collapse toward a simple “good/bad” judgment. These results provide evidence that emesis caused by LiCl modulates network activity in GC and plays an important role in shaping the cortical taste responses that are necessary for learning and decision-making, and further expand our insight into the state-dependence of sensory coding.

## Materials and Methods

### Subjects

Female Long–Evans rats (*n* = 26 250–300 g at time of surgery; Charles River Laboratory, Raleigh, NC) that were naïve to tastants served as subjects in this study. Animals were maintained on a 12h light/dark schedule, with experiments performed in the light portion of the cycle from 700 – 1100am. Rats had *ad libitum* access to food and water. All methods comply with the Brandeis University Institutional Animal Care and Use Committee (IACUC) guidelines.

### Apparatus

Neural recordings were made in a custom Faraday cage (6 × 24 × 33 cm) connected to a PC and Raspberry Pi computer (Model 3B). The Pi controlled opening time and duration of solenoid taste delivery valves. The PC controlled and saved electrophysiological recordings taken from electrode bundles *via* connections to an Intan data acquisition system (RHD2000 Evaluation System and Amplifier Boards; Intan Technologies, LLC, LA). Each bundle consisted of 32 microwires (0.0381mm formvar-coated nichrome wire; AM Systems) glued to a custom-made interface board (San Francisco Circuits) and soldered to a 32-channel Omnetics connector, which was fixed to a customized drive [29].

### Surgery

Rats were anesthetized using an intraperitoneal (IP) injection of a ketamine/xylazine mix (1mL ketamine, 0.05mL xylazine/kg body weight). Maintenance of anesthesia followed 1/3 induction dose every 1.25 h. The anesthetized rat was placed in a stereotaxic frame (David Kopf Instruments; Tujunga, CA), its scalp excised, and holes bored in its skull for the insertion of self-tapping ground screws and electrode bundles. The bundles were painted with a lipophilic membrane stain (ThermoFisher Scientific, Indianapolis, IN) and inserted 0.5 mm above gustatory cortex (GC; coordinates: AP +1.4 mm, ML ±5.0 mm, DV -4.5 mm from dura). Assemblies were cemented to the skull, along with two intraoral cannulae (IOCs; flexible plastic tubing inserted close to the tongue in the cheek and extending upward to the top of the skull) using dental acrylic [26]. The rat’s body temperature was monitored and maintained at ∼37° C by a heating pad throughout the duration of the surgery. Rats were given 6d to recover from the surgery.

### Water restriction

Mild water restriction (*ad libitum* access to 20mL at 4PM every day, 6 hours after experimental sessions), started 6d following surgery, ensured engagement in the task. Daily records were kept of weight and water intake to ensure rats did not fall below 85% of pre-surgery weight.

### Experimental Design

On the second day of water restriction, rats began 2 days of habituation to liquid delivered directly to the tongue via IOC, with 60 and 120 30-µL infusions of water delivered each day, respectively. For Experiment 1, on the morning following the 2^nd^ such session, *in vivo* recording sessions commenced: rats received a subcutaneous (sc; dose 0.50% of body weight) injection of isotonic Saline followed 20-min later by an injection of either 0.15M LiCl (saline-LiCl; **Fig. 1A**) or a second Saline injection (saline-saline); following each injection, neural and behavioral data were collected—20min of “passive” recording followed by 120 taste delivery trials in which one of four gustatory stimuli (0.1M sodium chloride [NaCl], 0.3M Sucrose, 0.1M Citric Acid [CA] and 0.01M Quinine-HCl [QHCl]) was delivered via IOC in a pseudo-random order. Stimuli and concentrations were chosen to ensure a range of distinct taste identities and palatabilities, and to maximize comparability to our [30-33] and others’ [34, 35] studies. A second testing session, identical to the 1^st^ with the exception that only a single sc. injection of Saline (0.50% body weight) was administered 20-min prior to taste delivery, was given 48hrs after the first testing session (**Fig. 1A**). To test whether any observed LiCl/NaCl differences were not order effects, a subset of animals (N = 5) were subjected to the same experimental protocol with the exception of the order of injections being reversed (two Saline injections on the 1^st^ day, LiCl injection on the 2^nd^). Our analyses revealed no difference between the orders (results not shown).

**Figure 1:**
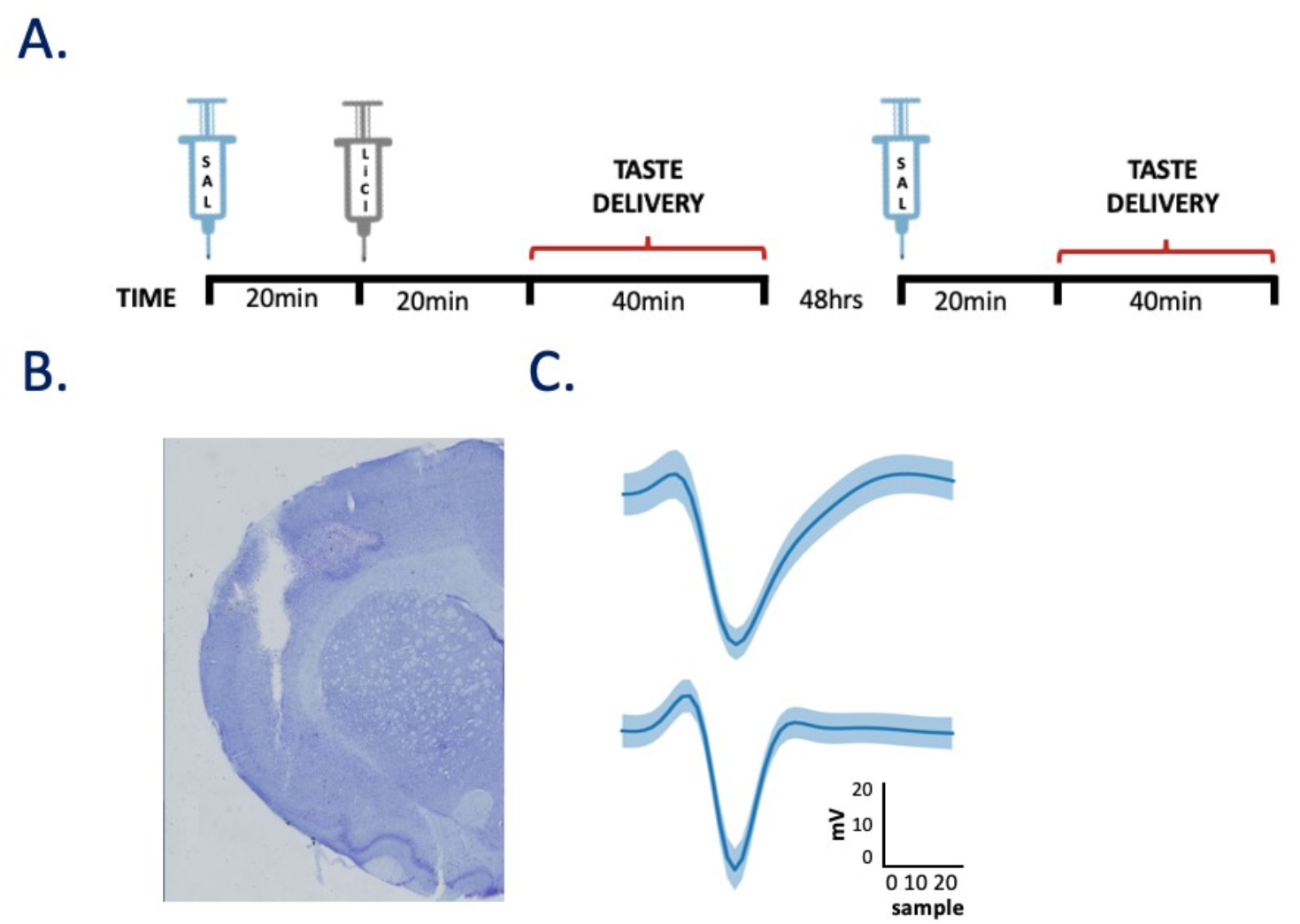
General sickness induction protocol and histological verification of recording location in GC. (A) On Day 1, rats received a single injection of Saline followed by an injection of 0.15M LiCl. Spontaneous GC activity was recorded for 20 min following each injection, after which tastes were delivered *via* IOC across a ∼40min session. On Day 2, only a single saline injection was administered. (B) A coronal slice from one rat stained with Cresyl violet. The track is an electrode bundle lesion, with the end of the lesion marking the final resting location of the wires. (C) The average waveform of representative regular [top] and fast-spiking [bottom] single neurons’ action potentials, with confidence intervals.

For Experiment 2, An additional 3 taste-naive animals were subjected to a single recording session identical to session 1 above, in which the injection of LiCl was immediately followed by taste deliveries *via* the IOC.

### Behavioral analyses

The assessment of lethargy provided a basic measure of sickness [2, 28]. Three cameras (positioned below, above, and diagonally within the behavioral chamber (see Apparatus) captured mobility in a separate subset of animals implanted only with IOCs (N = 5). Each video was manually scored for lateral (grid-line crossings) and vertical (rears; two forelimb paws off the ground for >0.5s) movements by a trained, but blind, observer. Such events have been shown to be sensitive measure of welfare in rodents—healthy, non-anxious mice/rats will perform lateral and vertical movements as a means to explore their environment [19, 36]. Given the small size of the recording chamber, the rats had limited ability to move laterally, which led us to focus our analyses primarily on vertical movements. Healthy/sick differences in frequency and duration of these movements were analyzed and fit with sigmoid functions so that we could determine whether and when LiCl-treated rats became ill.

### Histology

At the completion of the experiment, rats were deeply anesthetized with ketamine/xylazine (120:15 mg/kg, IP) and then perfused transcardially with physiological saline followed by 10% formalin. The brains were extracted and stored in a 10% formalin / 30% sucrose solution for at least 3 days before staining, after which they were frozen and sliced on a sliding microtome (Leica SM2010R, Leica Microsystems; thickness 60 µm). Slices were mounted, and sections were stained with a Nissl stain (ThermoFisher Scientific, Indianapolis, IN) to evaluate cannulae and electrode tracks respectively *via* inspection of fluorescence on a light microscope.

### Electrophysiology methods and analyses

Differential recordings from the micro-electrodes were sampled at 30 kHz using a 32-channel analog-to-digital converter chip (RHD2132) from Intan Technologies. The signals were digitalized online at the head stage/amplifier and saved to the hard drive of the PC. The collected recordings were filtered for either single-neuron action potential isolation (300-3000Hz bandpass) or local field potentials (LFP, 1-300Hz lowpass), and analyzed off-line.

Putative single-neuron waveforms (3:1 signal-to-noise ratio) were sorted using a semi-supervised methodology: potential action potentials were grouped into clusters by a Gaussian Mixture Model (GMM), which were then refined manually (to enhance conservatism) by the experimenters (for details see [29]). A total of 253 GC neurons were isolated across 12 sessions (2 sessions/animal) from six saline-LiCl group rats, and 190 GC neurons were isolated from six Control group rats.

LFPs were extracted only from channels containing isolated single-neuron data, in order to avoid potential signal artifact arising from noisy/broken channels. Data were averaged across electrodes within and across session for each animal, and analyzed across the 1-20Hz frequency bands, with focus on the µ/α bands (7-11Hz, [37-39] in one-minute time bins. After normalizing (across the entire session) each animal’s µ power to between 0 and 1, the LiCl vectors were subtracted from the Saline vectors to reveal changes in µ caused by illness, and an ANOVA was used to compare groups. Differences were considered significant only if 3 or more consecutive time bins reached a *p* < 0.05 criterion.

#### Change Point Analysis

To determine the position of a changepoint in GC µ power, we used a custom model (see fig x below) implemented in the PyMC3 probabilistic programming package [40], with parameter estimation performed using Markov-Chain Monte Carlo sampling (see details below). The LFP power timeseries were z-scored, and then modeled using two normal distributions; the model attempts to detect a single changepoint (tau) in the mean value of the LFP power as the timeseries switches between distributions. Changepoints were determined for each animal independently.

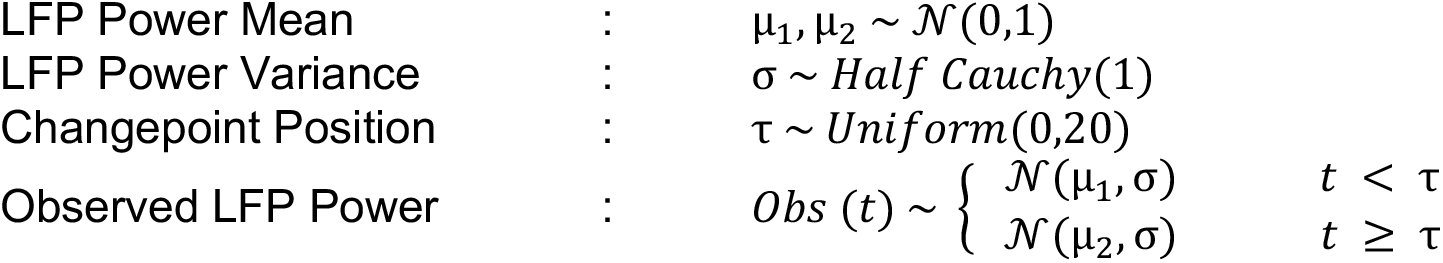

Since sampling returns a distribution over the parameter values explaining the data, distributions of tau with low variance (high peaks) indicate strong presence of a changepoint. To mark the position of these changepoints using the tau distribution in an unbiased manner, we compared the tau distributions (containing equal numbers of samples) of our timeseries (actual data) to those of 50 temporally shuffled timeseries (shuffled data) which, by definition, have no changepoints. Points in the actual data distribution higher than the 99th percentile of the shuffle distribution were marked. The average of these marked timepoints were taken to be the position of the changepoint for that timeseries.

#### Taste responsiveness

As in previous work [41-43], a neuron was considered taste responsive if taste-driven firing rates (0-2 s post-delivery) were significantly higher or lower (paired-sample *t-tests*) than pre-stimulus baseline activity (2s before taste delivery). Data were collapsed all four tastes, such that “responsivity” indicates purely that a GC neuron responds to taste delivery and reveals nothing about taste specificity.

To evaluate how LiCl changed the directionality (excitatory/inhibitory) of taste responses, we subjected these data to a Mann-Whitney U test, ignoring the first 300ms of post-taste activity (which is seldom chemosensory, see [42]). Chi-square tests assessed how LiCl impacted the number of significant responses, and paired-sampled *t-tests* (with Bonferroni correction for multiple comparisons) determined how LiCl impacted the magnitude of responses. The results from these tests revealed, at a macro level, LiCl significantly impacted neither the quantity nor magnitude of responses in a taste-specific manner. For this reason, we averaged taste responses (within condition) and applied a 2-way ANOVA to taste-responsive neurons, to test whether condition interacted with response direction on taste response magnitude. A subsequent series of 1-way ANOVAs was applied to each response direction to see how sickness impacted response magnitudes (collapsed across taste) across the taste processing epochs (Early: 100-350ms, Middle: 500-750ms, Late: 800-1050ms).

#### Taste specificity

To determine how LiCl impacted the taste specificity of GC responses, we used a standard linear discriminant analysis (LDA) classifier as brought to bear in previous studies [42, 44, 45]. This classifier tests the reliability with which a trial of taste evoked response can be identified among responses to other tastes. We binned neural responses into 250 ms bins and used a linear classifier with a leave-one-out (LOO) validation approach, calculating the prediction accuracy of the classifier averaged across each excluded trial for each time bin. Paired-sample *t-tests* (with Bonferroni correction) were performed for each post-taste delivery epoch (see above) to determine the impact of LiCl.

To assess changes of taste discriminability within a single session, the same LDA classifier was used, trained on the first five trials (per taste) and tested on later trials. Across-iteration averages were computed for each tested trial and the resulting means were binned into 5-trial blocks (∼6min/block). Blocks were normalized (between 0 and 1) across animals and results were subject to a repeated measures ANOVA and a series of paired T-tests (with Bonferroni correction).

#### Taste Palatability

Using our now-standard palatability-correlation analysis [31, 44, 46, 47], we evaluated the degree to which LiCl altered the amplitude of palatability-relatedness in late-epoch GC taste responses. Using a moving window (window size: 250ms, step: 25ms), we correlated firing rates with well-established palatability ranks (Sucrose > NaCl > CA > QHCl, [44, 48], and compared the magnitude of this correlation in healthy and sick rats.

We also computed what we call a “pure palatability index” (PPI), adapting methods from [49], testing the degree to which GC taste responses coded a simple “good vs bad” dichotomy by comparing the Euclidian distances between single neurons’ responses (normalized to -500ms pre-taste delivery) to tastes with similar palatability (i.e., Sucrose and NaCl, CA and QHCl) to those for tastes with different palatability (Sucrose and CA, Sucrose and QHCl, NaCl and CA, NaCl and QHCl). A lack of palatability-related information results in a PPI of 0 (because tastes of similar palatability and tastes of different palatabilities are equally different), and the more polarized the response into “good vs bad,” the more positive the PPI. To avoid artificially attenuating the effect via the inclusion of consecutively linked timepoints, the PPI was calculated from firing within the middle of our standardly used Epochs [42, 43]. The results were significance tested using a Wilcoxon sign-ranked test.

#### Stability of single-neuron waveforms across days

A subset of analyses required the stable tracking of neurons held across testing sessions—“held neurons.” To evaluate whether a neuron was “held,” we performed a spike shape analysis brought to bear in several previous studies [30, 32, 50, 51]. For the equations and clustering statistics used to isolate held units, refer to [32]. Implementing such metrics, we applied a conservative criterion (comparing each neuron’s waveforms from the 1^st^ third of the session to those of the last third) for each condition (saline versus LiCl) such that only neurons for which the between-session (non-parametric clustering statistic) value was less than the 95^th^ percentile value calculated from within-session, were counted as ‘held.’ Of the entire population of recorded GC neurons, a total of 55 (21.7%) were determined to be stably held across testing sessions.

#### Cluster detection in patterns of healthy-ill Response Differences

A clustering analysis was used to seek patterns in how illness impacted palatability-related firing in individual held neurons. A condition response was determined for each held unit (under each condition) by taking the average (minmax-normalized) pre (−750à-250ms) and post (250à750ms) -delivery firing rates (presented as a percentage of maximum responsiveness). We then calculated and plotted the distance and Cartesian direction between condition responses or each neurons’ taste response, yielding 220 (55 held units X 4 tastants) response differences (RDs)—measures of how illness changed the excitatory or inhibitory response to the taste. As a conservative estimate of RD likeness, we determined the number of clusters that best fit a GMM probability distribution (calculating the Bayesian information criterion) as done previously [52]. Classifications were considered valid only if they fell within the 95% confidence interval from the centroid of each respective cluster (see **Fig. 8C**), and we further constrained our analyses to neurons for which all RDs were classified to the same cluster.

#### Cluster-specific Taste Palatability

Using methods described in ‘Taste Palatability,’ we evaluated the effect of condition on palatability-related firing for each neuron’s response for every cluster. A non-parametric 2-way ANOVA was employed to indicate whether differences (Saline – LiCl) in hedonic-coding (Spearman *rho*^2^) differed across cluster and time during taste-processing.

## Results

### LiCl-induced illness changes GC LFPs just prior to changing behavior

We recorded spontaneous and taste-driven activity following subcutaneous injections of either LiCl or Saline (**Fig. 1A**); GC single-unit responses (**Fig. 1C**) and LFPs were acquired from a drivable bundle of 32 wires (**Fig. 1B**). Given that network function, measured in terms of spectral properties of LFPs, changes with even the most innocuous of body states (e.g. sleep versus wake, [53, 54]), and that single-neuron firing is altered in concert with these changes ([55, 56]), our investigation into the characterization of illness started with assessment of changes in GC LFPs. We focused on power in the mu/alpha (µ: 7-12Hz; [25]) range, as power in this frequency band has proven particularly sensitive to changes in even general states related to wakefulness and attention [20, 25, 26, 38, 57]. While the results described below were observed in frequency ranges above (β) and below (θ) µ, they were centered on and largest in the µ range.

Save for the brief period immediately following handling and injections, the amplitude of µ in “spontaneous” activity remained relatively stable across the first half of the (20min) post-injection recording periods (**Fig. 2A**). Differences between the impact of saline and LiCl injections emerged in the second 10 min following injections. The precise nature of this change varied with individual (data not shown), in some cases (N =3) involving a reduction of µ power and in some cases an increase (N = 4), but a change point analysis (CPA) quantifying the time-points and likelihoods of change in mu power for each animal (across both experimental groups) revealed that LiCl-induced changes in GC µ power reached significance at around the same time in 5 out of 7 (see Methods) rats (**Fig. 2B, top**). Had no change occurred, or had the change been gradual/subtle following injection (as in saline-saline injected rats, **Fig. 2B, bottom**), the likelihood of change (*τ*; posterior distribution) would be spread out across time; instead, the analysis reveals that a change in GC µ power occurs between 10.89 and 17.07min post-injection, with the mean likelihood occurring at 15.11min.

**Figure 2:**
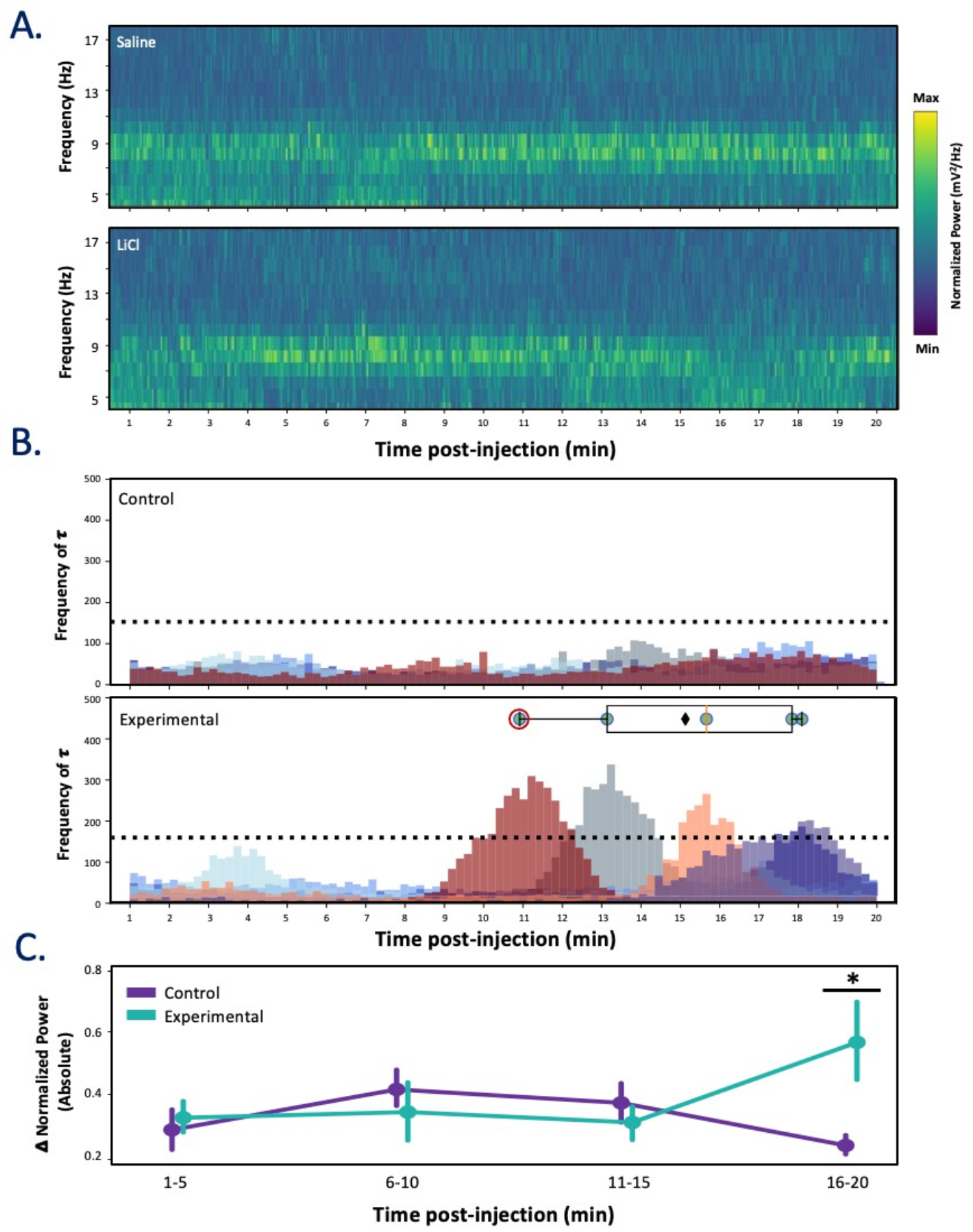
Sickness alters spontaneous GC Local Field Potentials 15-minutes post-injection. **(A)** representative spectrograms from one animal showing spontaneous activity post-Saline [top] and -LiCl [bottom] injection. With respect to Saline, LiCl reduces power in mu beginning around 11min post-injection. **(B)** The posterior distributions for a single changepoint model identifies the probabilities of a change (shown as frequency of *τ*) in mu power (each color represents distribution for single animal) for Control (top) and Experimental (bottom) groups. Dashed line indicates the 99^th^ percentile from temporally shuffled data (see Methods) where the overlaid boxplot depicts the range in peak likelihood of change in mu power for distributions with values higher than 99^th^ percentile relative to shuffled data; each dot represents a single animal (animal from panel A encircled in red). Box extends from lower to upper quartiles (red line: median, black diamond: mean) of likelihood. The mean onset of the change in mu power occurs at 15.11min, with all changepoints occurring after 10 min post-injection. **(C)** The absolute difference (Saline – LiCl) in mu for Control (purple) and Experimental (teal) groups reveals a significant interaction effect on change in mu power between group and quartile (*p <* 0.05, two-way ANOVA), with LiCl changing mu power significantly within the 4^th^ quartile (* *p* < 0.05; one-way ANOVA). Error bars represent SEMs.

An analysis of the group data confirms these rat-by-rat results, demonstrating that absolute changes in µ power following LiCl injection emerged late in the 20-min post-injection recording session (compared to saline-saline animals, **Fig. 2C**). A two-way ANOVA on these data revealed a significant interaction between the group and quartile (*F*(3, 33) = 3.62, *p* = 0.023); subsequent testing confirmed that the difference reached significance only in the 16-20 min post-injection bin (*F*(1, 11) = 5.14, *p* = 0.04). While one could speculate that illness’ effect on GC µ power merely represents changes in firing rates (FRs) across time, a two-way ANOVA revealed a there was no significant interaction between the experimental group (saline-LiCl versus saline-saline) and quartile on change in FR (*F*(3, 33) = 2.39, *p* = 0.09); this suggests that our observed effect on µ is not simply a gating of firing rate direction, but rather a matter of synchrony being modulated.

Overall, these results accord well with those of previous studies, which have demonstrated: 1) that changes in LFP activity are hallmarks of the onsets of cortical state changes, regardless of the specific directionality of the changes [58-61]; 2) that these changes are typically centered on the µ band [62]; and 3) that sickness-related behaviors such as immobility emerge at approximately this timepoint following LiCl injections [19, 36, 63] [64-66].

To independently confirm this last point, we compared our LFP data to (independently collected and blindly coded) video recordings of illness-related behaviors, predicting that the above-described changes in network activity would foretell the time at which changes in mobility appeared following LiCl injections. We focused our measurement of mobility on rearing events (the small recording chamber limited the rats’ ability to move laterally) in which the animal lifted both forepaws off the floor simultaneously without proceeding to grooming [67, 68]. We specifically hypothesized that reduction of such rearing events, which has been linked to LiCl-induced illness in previously established findings [66], would occur at around the time that we observed changes in cortical LFP µ power (**Fig. 3A**).

**Figure 3:**
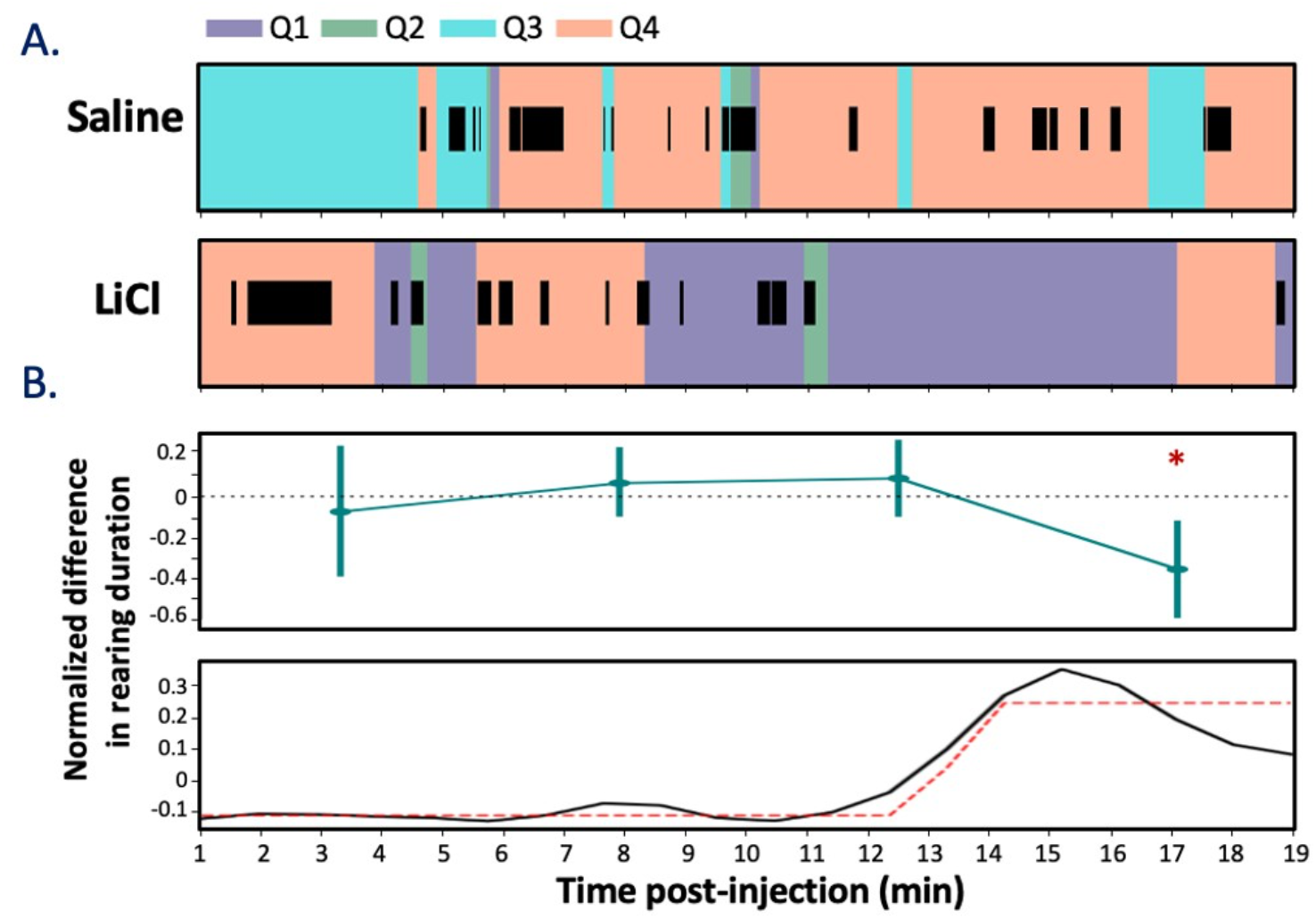
Sickness reduces rearing ∼12min after LiCl injection. **(A)** An example rat’s movement through time around the cage space (color-coding shows quadrant position, Q1-4) overlain with rearing (black bars) movement durations after Saline [top] and LiCl injections [bottom]. **(B)** The average normalized LiCl – Saline difference in rearing duration across animals reveals [top] a significant reduction occurring in the 4^th^ quarter of the session (* p < 0.05, Wilcoxon-Rank sum). A more fine-grained analysis [bottom] suggests that the change in behavior (black line) begins at 12min post-injection; the dashed red line illustrates the sigmoidal fit (*r*^*2*^ *=* 0.72). Error bars represent SEMs.

In fact, the durations of rearing events did decline following LiCl injection. With the 20-min recording session \again binned into 5min quartiles (**Fig. 3B, top**; large bins were used to ensure sufficient power), a two-way repeated measures ANOVA revealed a significant difference in rearing duration emerging post-injection (*F*(3, 182) = 3.14, *p* < 0.05). This change, like the change in cortical LFP power, became significant (according to post-hoc Wilcoxon Signed-rank tests) in the 15-20min post injection bin (W = 158, Z = 0.65, *p* = 0.024, *r* = 0.50), compared to saline-saline rats. A more sensitive sigmoidal curve fit to the normalized LiCl-saline difference in rearing duration (**Fig 3B, bottom**) revealed the appearance of this illness-related change in behavior (the asymptote minima of fit; *r*^*2*^ *=* 0.72) to align well with the above-noted change in GC µ power (compare to **Fig. 2, top**), suggesting that the onset of LiCl-induced illness is reflected in the function of the GC network; the fact that the **Fig. 2** and **Fig. 3** data were collected in separate groups of rats only increases the conservatism of this interpretation (see Discussion).

### Illness causes taste responses to resemble a good-bad distinction

As an initial look at how illness impacts taste-evoked spiking activity of GC neurons, we evaluated the quantity and magnitudes of responses elicited by tastes delivered directly into the mouth *via* IOC. Paired-sampled *t-tests* (with Bonferroni correction for multiple comparisons) revealed that LiCl did not significantly impact the magnitude of responses in a taste-specific manner (*ps* <0.05). For this reason, we averaged taste responses (within condition) and a subsequent chi-square test of independence revealed a significant relationship between condition and direction of response on neuron count *X*^*2*^ (1, 2) = 9.21, *p* = 0.009). LiCl impacted the number of neurons that had inhibitory or unchanged responses (**Fig. 4A**).

**Figure 4:**
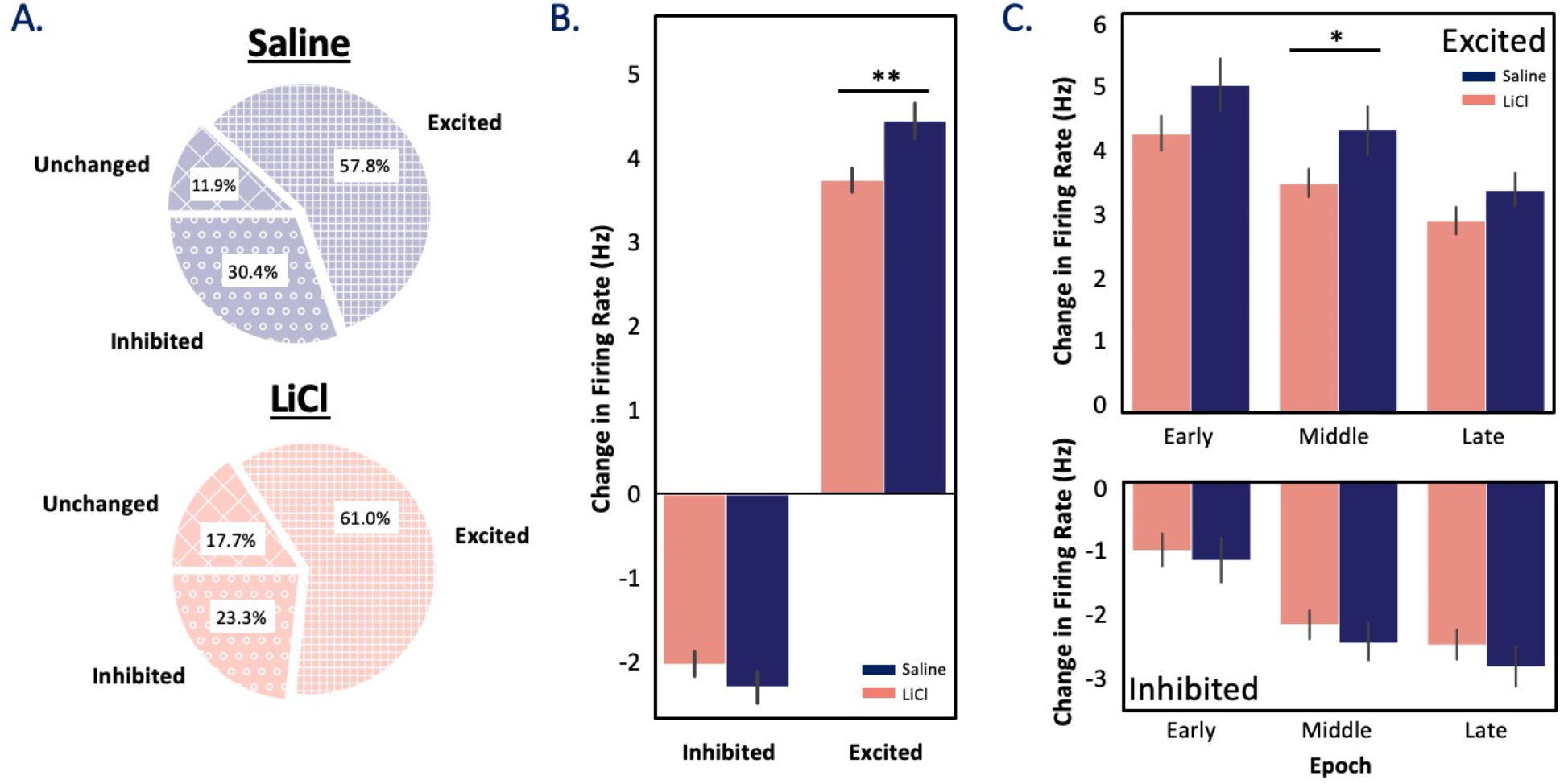
Sickness reduces magnitudes of taste responsiveness. **(A)** LiCl-induced sickness had a significant impact on the relationship between condition and response direction; chi-square test on each neuron count, *X*^*2*^ (1, 2) = 9.21, *p* = 0.009). **(B)** The magnitudes of these responses revealed an interaction between condition and the direction of taste response – neurons with excitatory responses experienced a significant reduction in this response due to illness (** *p* < 0.01; two-way ANOVA with Tukey post-hoc). **(C)** Examined across time (epoch-wise), the significant differences between conditions for excitatory responses can be seen to emerge within the identity epoch of taste processing (* *p* < 0.05, one-way ANOVA). Error bars represent SEMs.

The magnitudes of these responses were impacted by illness, however: a two-way ANOVA revealed a significant interaction between the condition and taste response direction (*F*(1, 3191) = 6.05, *p* = 0.014), with a Tukey post-hoc comparison showing the main effect of illness occurred for excitatory responses (*p* = 0.003, **Fig. 4B**). Following a series of one-way ANOVAs applied to each taste epoch, this impact of LiCl on excitatory taste responsiveness proved maximal during the “middle epoch” of the taste responses (**Fig 4C**, *F*(1, 735) = 4.14, *p* = 0.042))— the period known to contain firing that codes the identity of taste stimuli in GC [42, 43]. This result suggests that illness perturbs the coding of taste discriminability.

To test this hypothesis, we performed a linear discriminant analysis (LDA), quantifying the reliability with which GC responses to one taste could be differentiated from responses to other tastes. As revealed in **Fig. 5**, our hypothesis was borne out: LDA allowed us to correctly identify administered tastes from middle-epoch responses on 40-50% of the trials in healthy sessions (well above chance, which = 25%), despite using a small time-bin sliding window analysis that left the analysis at the mercy of unsmoothed trial-to-trial variability. The distinctiveness (i.e., classifiability) of responses was significantly diminished following systemic LiCl administration, however, with the decrement becoming significant in the “Middle Epoch” and continuing into the “Late Epoch” (Mann–Whitney *U* (one-tailed) = 1952, *μ* = 34.06%/43.77%, SD = 11.09%/14.58%, *p* = 0.0003; *U* = 2045, *μ* = 39.78%/47.83%, SD = 15.72%/14.64%, *p* = 0.001, respectively). Taste responses carry less identity-related information during illness.

**Figure 5:**
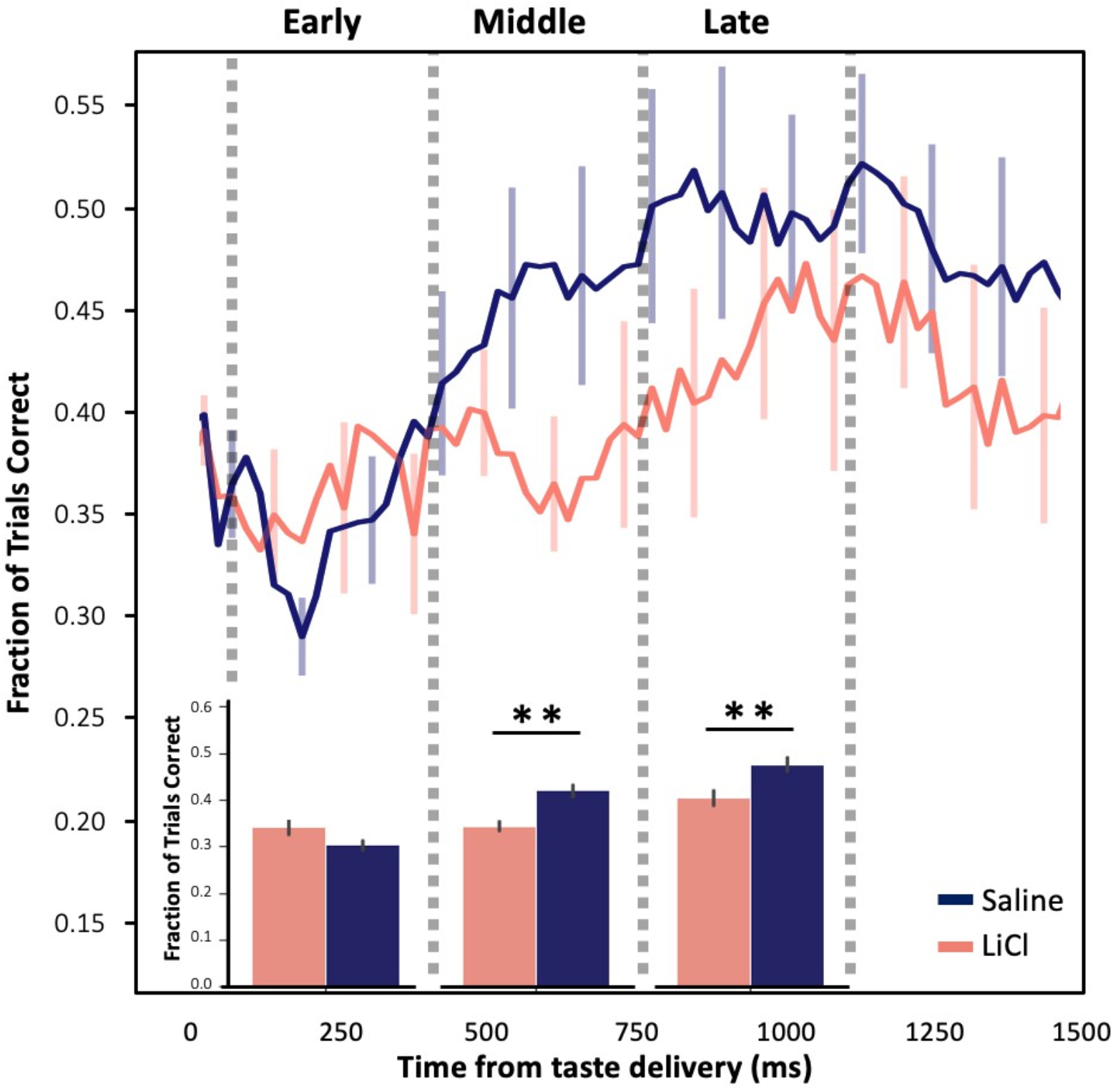
Sickness reduces taste discriminability in neural ensembles. LiCl-induced illness decreases taste discriminability in GC neurons in discrete time windows. **(top)** Time series of Linear Discriminant Analysis (LDA) shows the fraction of trials correct is reduced when an animal is sick (coral) in comparison to when they are healthy (purple). **(bottom)** These differences only become significant within the middle (400-700ms) and late (800-1100ms) epochs post taste-delivery (** *p <* 0.01; pairwise T-test with Bonferroni correction). Error bars represent SEMs.

The fact that this decrement in discriminability extends into the “Late” (i.e., palatability) epoch led us to ask whether LiCl-induced illness might also change the palatability-relatedness of coding as well (a feature that is known to emerge relatively late in GC taste responses, see [42, 43]). We began with the simple hypothesis that reduced discriminability should imply reduced palatability-relatedness, testing this hypothesis in the standard manner [31, 44, 46, 47], i.e., performing moving window correlations between firing rates and the known canonical palatability ranks of the administered taste stimuli.

As has been observed many times previously, palatability-related information in our GC taste responses climbed (regardless of condition) across the period leading into the Late epoch (**Fig. 6A**; note that we currently have no explanation for the small, low-magnitude, but significant difference in palatability-related firing in the earliest responses, but see Discussion). The climb calculated following saline and LiCl injections diverged significantly, however, as the Late Epoch was reached – peak correlations following LiCl injections were higher than those observed in saline condition within this epoch (H(1) = 8.68, *p* = 0.003). This means that our initial hypothesis was disconfirmed: whereas illness reduces Middle epoch taste response discriminability, it increases Late epoch taste response palatability-related content.

**Figure 6:**
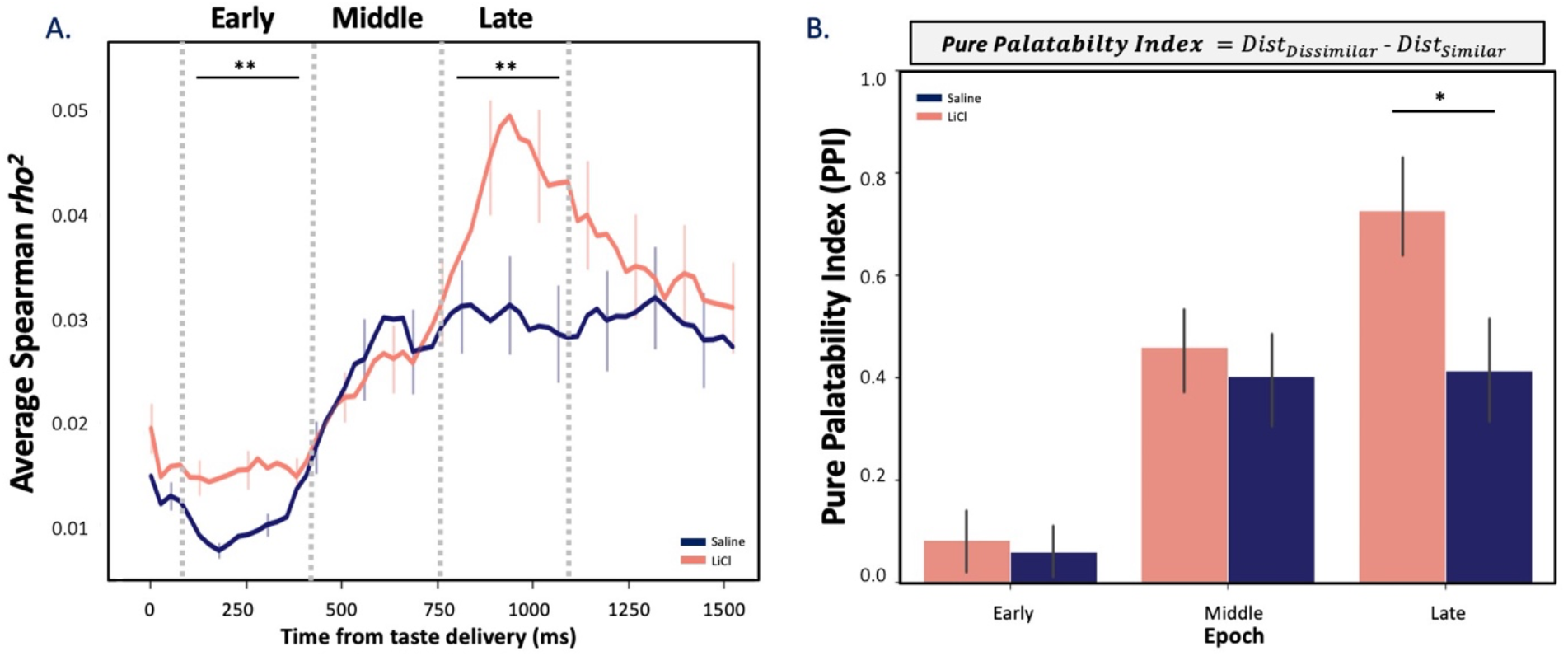
Sickness increases the palatability-relatedness of GC taste responses. **(A)** Average correlations (Spearman rank-correlation coefficient) between firing rates and known palatability grew stronger as responses neared the Late epoch (as noted many times previously). This correlation rise reached a higher peak following LiCl injection than saline (Kruskal-Wallis H-test, ** *p <* 0.01). **(B)** The sickness-induced reduction of taste discriminability, juxtaposed with this sickness-induced enhancement of palatability-relatedness, suggests that illness selectively decreases the distinctiveness of tastes with similar palatability. As predicted, a “pure palatability index” (PPI), which evaluates this possibility (see inlaid equation), peaks a a larger Late epoch value following LiCl injection than following saline (unpaired 2-sample Wilcoxon test; **\*** *p* < 0.05), reflecting sickness-induced polarization of coding. Error bars represent SEMs.

The above results beg the question “how can illness reduce information pertaining to identity while enhancing the palatability-relatedness of the same taste responses?” We hypothesized that the reconciliation of these seemingly contradictory results requires that illness causes GC coding to reduce toward a simple “good vs bad” judgment, preferentially decreasing the differences between the coding of tastes with similar palatabilities—making Sucrose and NaCl (the palatable tastes) responses more similar and making quinine and citric acid (the aversive tastes) more similar—thus leaving GC responses closer to a simplistic, “pure” code of whether or not a taste is palatable. We tested this hypothesis by quantifying the distance in Euclidean space between neural responses for all pairs of similar (e.g., Sucrose and NaCl) and dissimilar (e.g., Sucrose and QHCl) taste stimuli. A “pure palatability index” (PPI; difference in distance between the similar and dissimilar taste pairs) value of zero would indicate a lack of any sort of palatability-related responsiveness, while a PPI of one would indicate that GC neural responses are determined purely on the basis of whether the taste in question is pleasant or aversive.

**Figure 6B** shows the results of this analysis. One-sample T-tests revealed that the PPI reaches significance only after the Early epoch in both conditions (*ps* < 0.01), and a subsequent Wilcoxon Signed-rank test (performed because the data differed significantly from normal, *p* >0.05) revealed that LiCl-induced illness significantly enhances the PPI in the Late epoch (W = 68379, Z = 0.54, *p* = 0.037, *r* = 0.09), reflecting enhanced polarization of the coding of palatable and aversive tastes. Thus, our hypothesis was confirmed: LiCl shapes the coding of taste hedonics by enhancing the polarization between similar and dissimilar tastes in the Late epoch GC taste responses, and thereby simultaneously increases the overall palatability correlation and reduces the average discriminability of the responses.

### Illness-related changes in taste coding occur in single neurons

The above results suggest that ensembles of GC neurons assayed 10-30min after illness induction code tastes differently than ensembles of GC neurons assayed in healthy rats. The implication of these results is that coding in individual neurons changes as sickness emerges. To directly test this implication, we collected data using a modified experimental protocol in which we administered (and acquired spiking responses for n = 74 GC neurons to) tastes immediately after the injection of LiCl (**Fig. 7A**). As already established, illness-related behaviors and changes in GC µ power emerge between 10 and 17 minutes after LiCl injection (**Fig. 2 and 3**); we therefore hypothesized that the above-described coding changes would emerge within single ensembles following this time point—that coding in trials delivered before sickness (“Pre”) would differ from that in trials delivered after sickness onset (“Post”; **Fig. 7A**).

**Figure 7:**
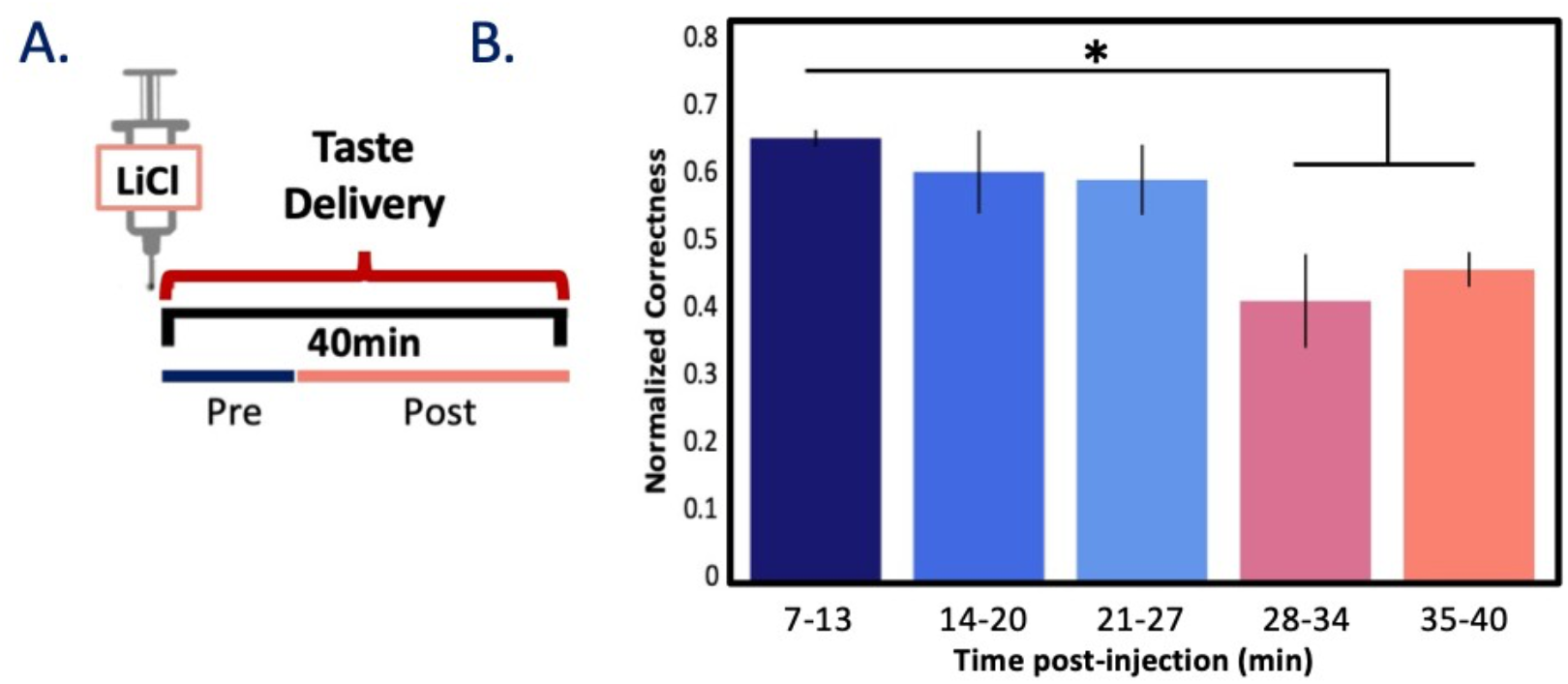
Reduction in taste discriminability emerges with sickness within session. **(A)** A schematic of the 2^nd^ testing protocol, in which an injection of 0.15M LiCl was immediately followed by delivery of tastants *via* IOC for ∼40min. **(B)** Bar plot shows the percent of trials correctly predicted using a Linear Discriminant Analysis (LDA; trained on the first 5 trials) from GC responses against the tested trial. Across N = 3 animals (n = 56 taste responsive neurons), taste discrimination is significantly reduced post-sickness induction. * *p <* 0.05; paired T-test with Bonferroni correction. Error bars represent SEMs.

We again used LDA to test whether and when taste response coding changed, by testing the similarity of the first five ‘healthy’ trials of each tastant to each subsequent taste delivery (binned; 5 trials/taste/bin). As expected, a repeated measures ANOVA performed between bins (bin =6min, ∼5trials/min) on taste responsive neurons (n = 56) revealed the classifiability of tastes to decrease significantly as rats became ill (*F*(4, 8) = 5.42, *p* = 0.02, *np*^2^ = 0.64), with this disruption becoming significant ∼30mins into the session (**Fig. 7B**, *<u>ps</u>*<u>< 0.05</u>). This result is a good match for the above-described changes in GC *μ* power and illness-related behaviors (**Figs 2 – 3**). While one could speculate that this effect is driven simply by time (as opposed to illness), this explanation is rendered unlikely by previous results demonstrating that, in the absence of illness, within-session taste discriminability does not vary significantly across much longer time spans [26]. Clearly, taste responses change with the onset of illness.

We went on to more closely examine the precise nature of these changes in the subsample of our recorded neurons that were held across sessions (see Methods for criteria)— neurons in which we were able to assay responses after both LiCl and Saline injections. For each of 55 held neurons, we first examined session-specific firing rates for the 500ms before and after taste delivery. In a scatterplot of these basic firing properties from both sessions (**Fig 8A)**, pre-stimulus firing rates fall generally along the diagonal, demonstrating that spontaneous firing rates do not differ strongly between condition. Comparisons of the taste responses, meanwhile, suggest three distinct “types” of illness impact (accentuated with distinct colors attached to different patterns of difference). Subsequent analyses confirmed these appearances (**Fig 8B-C**), revealing three distinct and discrete clusters of taste-response changes wrought by sickness: 1) in some neurons, taste responses were excitatory in saline sessions but inhibitory during illness; 2) in others, the reverse was true; and 3) in a third group of neurons, responses were excitatory in both states, but these excitatory responses were generally less strong during illness.

**Figure 8:**
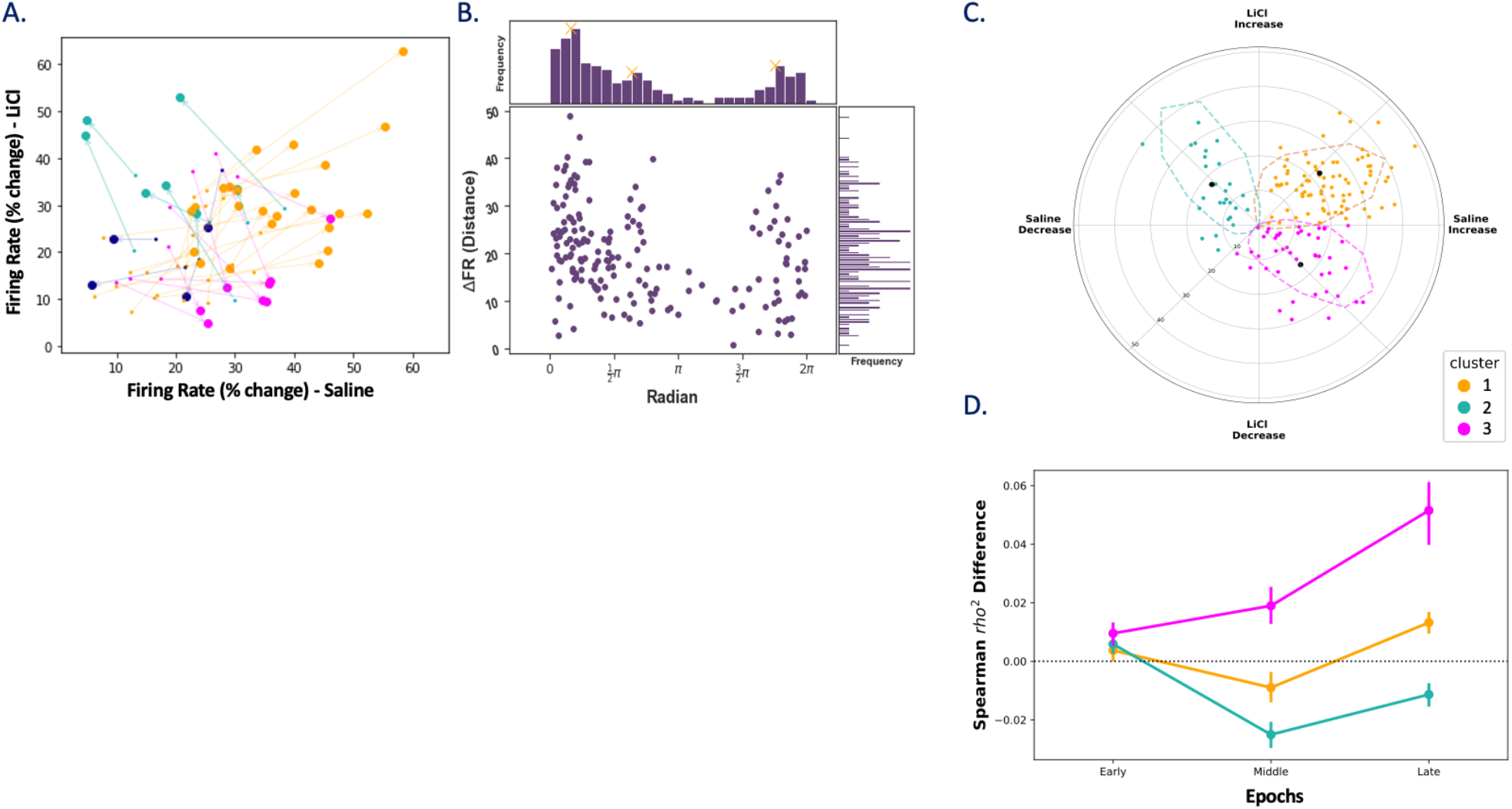
Neurons held across post-LiCl and post-saline tasting sessions produce distinct “types” of taste response profiles. **(A)** normalized firing rate trajectories for single neurons before (“pre,” small circles) and after (“post,” large circles) taste delivery (here, to citric acid) reveal condition-specific (sickness – y-axis, healthy – x-axis) taste response properties. Responses are colored with respect to angle of the response trajectory – 0-90°, 90-180°, 180-270°, and 270-360° are yellow, teal, magenta, and blue respectively – which in turn represent the nature (excitatory, inhibitory) of responses in each condition. **(B)** A population response histogram depicting each neuron’s taste response (change from pre taste) as a function of response direction confirms the presence of three unique clusters of sickness impact (peaks indicated with X). **(C)** A Gaussian Mixed Model (GMM) was used to cluster cells based on response magnitude and trajectory revealing distinct cluster responses lie in a polar plane (dashed lines denote 95% confidence interval of cluster classification with respect to black filled centroids). **(D)** Cluster three (magenta) neurons reliably and selectively experience a sickness-induced increase in palatability-relatedness in the Late epoch (nonparametric 2way-repeated measures; *** *p* <0.001). Vertical lines represent SEMs.

These three clusters of neurons, which were sorted without regard to the types of information within the taste response, nonetheless differed with regard to how illness impacted palatability-relatedness dynamics of the taste responses. **Figure 8C** shows the result of the application of a three-component gaussian mixture model (GMM) to all responses; a 3way non-parametric mixed ANOVA was ran and revealed a significant interaction between cluster, epoch, and condition, *F*(4, 478) = 4.80, *p* = 0.001, indicating that the relationship between cluster (whether the neurons were in cluster 1, 2, or 3) and the coding of palatability within epoch was significantly different across the Saline and LiCl conditions. Following a series of paired T-Tests, these data revealed that the Late-epoch enhancement of palatability-related firing was primarily a function of neurons for which normally excitatory responses were turned into inhibitory responses by LiCl-induced illness (*F*(1, 130) = 33.47, *p* = 5.12e-8; **Figure 8D**). This result both confirms the validity of the separation of neuron types and reveals that this separation is related to the impact of illness state on palatability processing.

## Discussion

Nutritional requirements, environmental conditions, and experience work together to guide consummatory behavior [2, 11, 13], in cooperation with an animal’s internal state (e.g., hunger, illness) [1]. This necessitates the study of how physiological states impact perception, as this relationship intrinsically shapes the likelihood of survival. The fact that neurons are embedded in networks, and that the states of such networks can be indexed in terms of EEG power (particularly in θ and µ frequency ranges [25, 26, 56], motivated our decision to characterize changes in LFP activity within GC as an animal shifts from a healthy state into one of emesis, and to then relate this shift to changes in the dynamics of taste coding. This analysis allows us to show that LiCl causes a modulation of µ power at approximately 15mins post-injection—a result consistent with human [69] and rat data [70]. In both the literature and our current data, the latency of these changes in µ are strongly correlated with the appearance of illness-related behaviors [70]. We show that rearing durations, a measure of healthy exploratory behavior [2, 64, 65, 70-72], drop starting at approximately 12min post-LiCl injection. This finding is consistent with decades of work showing that poisoning-related behaviors, and specifically those brought on by systemic nausea, emerge (depending on dose) at roughly 10min post administration [2-4, 70]. In our hands, the emergence of this illness-induced behavioral change lags closely (∼55sec) behind the change in GC µ power induced by the same LiCl dose, a fact that is all the more striking given that the neural and behavioral datasets were collected independently. These neural changes appear to be part of the process whereby an animal experiences illness.

The change in GC µ power is phasic, meaning that it is not simply the case that health connotes one µ amplitude and illness a different µ amplitude. This concept is not novel—while body states *can* be indexed in terms of LFP power fluctuations, behavioral states (e.g. sleep/wake) and muscle movements do not always correspond with a particular amplitude of field potential discharge [73], nor do significant changes in LFP power necessarily imply an observable change in body states [74]. Like many dynamical systems, the cortex experiences transient periods of instability at the time of state changes; we postulate that the observed transient impact of LiCl administration on GC µ power signals the onset of an illness state, and that the relaxation to a lower-power regime shortly (∼7min) thereafter nonetheless leaves the network *changed* in a way that impacts the processing of internal and external stimuli. In this, GC is akin to an automobile engine: shifting from one gear to the next requires a brief transition from a steady state to a transient and then back to a steady state for optimal efficiency [75, 76].

As predicted, illness impacts taste responses (despite the spectral transient having subsided). At the most general level, it reduces these responses’ magnitudes, with the reduction being most notable in the epoch (500-750ms post taste-delivery) known to code the identity content of taste stimuli. It makes sense to suppose that illness disrupted the coding efficacy of taste identity, decreasing the discriminability of GC taste responses. The finding that sickness reduced the magnitude of identity information in GC neural responses in both the taste identity epoch and the late (palatability) epoch, meanwhile, motivated our hypothesis that illness likely disrupts the coding of taste hedonics. Our test of this hypothesis showed, however, that despite reducing the overall discriminability of GC taste responses, sickness in fact significantly enhanced palatability-relatedness in the late epoch of these responses. We also observed a small but significant illness-related increase in palatability coding in the early epoch. While an explanation for this unexpected result awaits further experimentation, we would speculate, based on work showing that expectation of stimulus availability can reduce that latency of gustatory coding [35], that illness also enhances the ‘readiness’ of cortex, reducing the latency of taste-specific coding; since illness biases GC toward palatability-related coding (see below), it could very well be palatability coding that appears early when the animal is ill.

The fact that illness reduced the discriminability of GC taste coding while simultaneously enhancing the palatability-relatedness of those codes motivated a follow-up question: how could illness reduce taste identity information while enhancing taste palatability content? In order for these two results to be simultaneously true, we reasoned that illness must move the system in the direction of eschewing taste-specific information in favor of a pure “good vs bad” code – enhancing palatability-relatedness by specifically reducing differences between responses to tastes with distinct tastes but similar palatabilities, and thereby reducing overall discriminability. In essence, illness enhances palatability processing in GC which in turn attenuates identity processing. This is precisely what we found: illness changes GC processing of tastes in a manner that perhaps best suits the needs of the sick animal, de-emphasizing what the taste is in favor of whether it is helpful/harmful.

These findings, and interpretation thereof, may at first blush appear to be in conflict with previous studies examining behavioral changes following LiCl-induced illness [77-79], which suggest that sickness has no effect on palatability (e.g., as measured *via* evaluation of taste reactivity to a single taste [77]). In fact, however, our findings regarding sickness polarizing palatability processing are largely consistent with these results, in that they do not suggest wholesale changes in palatability of individual tastes, or even changes in the order of preference. Rather, we argue that illness simplifies palatability processing, making aversive tastes more similarly aversive and palatable tastes more similarly palatable.

These results suggest, but do not prove, that single neurons change their coding as a function of illness. It is always possible that neurons coding taste identity and palatability during illness represent a distinct set of neurons from those coding these properties in healthy rats, and that the differences in overall sample responses reflected the addition of this new set of “illness-only” responses. In the first test of the hypothesis that illness changes the processing of taste stimuli in individual neural ensembles, we showed that taste discriminability is significantly changed as an animal enters the emetic state. And since the small number of trials available in this analysis rendered it impossible to reliably compare palatability correlations, we performed a second analysis in which we leveraged our ability to hold a subset of our single neurons across both testing sessions. This analysis revealed that illness enhances palatability-relatedness in an epoch-specific manner but went beyond this to reveal three distinct subpopulations of illness-related response profiles.

We cannot, as of yet, provide a definitive explanation of this functional dissociation of GC neurons, but we can speculate as to several mechanisms that could explain our results. The most parsimonious of these explanations might be that the clusters come from either functionally or physiologically distinct populations of cells which in turn code specific features about the state of the animal (and thus responses to external stimuli) in unique, yet beneficial ways. However, cross-correlational analysis (methods adapted from Li et al [2013]; results not shown) failed to reveal differences in the strength of the functional connectivity between (nor within) the individual clusters, which would have been expected with such an explanation [48]. Furthermore, analysis of wave shapes and basal firing rates failed to reveal groups to be made up of different percentages of putative pyramidal and interneurons. While it is always risky to reach conclusions on the basis of a null result, particularly in small datasets, we find it unlikely that these clusters represent distinct types of GC neurons.

An alternative explanation for the observed differences in palatability coding across these putative clusters has to do with the possible sources of input to these neurons. It has recently been shown that specific Area Postrema (AP) neuron types are responsible for providing illness-related information to brain regions [18], and that these influences may reach GC by way of either the nucleus of the solitary tract [80] or other regions (e.g. amygdala) known to be impacted by the physiological state induced by LiCl [81]. It is possible that the palatability-coding differences between GC clusters is a downstream result of cell-type specific projections. Future work will examine this possibility.

Regardless of the answers to these questions, the work presented here demonstrates that illness, or at least that caused by LiCl, impacts cortical function in a far more refined manner than *via* a simple whole-sale disruption of activity. The effects of illness on GC activity are complex, but this complexity seems linked to an animal’s need, when ill, to simply tell good from bad. The fact that illness-related behaviors are observed to trail shortly behind changes in global activity in GC, accentuates the importance of assessing the welfare of an animal as it relates to both spontaneous and stimulus-driven activity.

## Acknowledgements

The authors would like to acknowledge and thank all members, past and current, of the Katz laboratory for their feedback and insightful comments. This work has been supported by National Institute on Deafness and Other Communication Disorders Grants R01-DC006666 & DC007703.

